# CoolBox: A flexible toolkit for visual analysis of genomics data

**DOI:** 10.1101/2021.04.15.439923

**Authors:** Weize Xu, Quan Zhong, Da Lin, Guoliang Li, Gang Cao

## Abstract

We developed CoolBox, an open source toolkit for visual analysis of genomics data, which is highly compatible with the Python ecosystem, easy to use and highly customizable with a well-designed user interface. It can be used in various visualization situations like a Swiss army knife, for example, to produce high-quality genome track plots or fetch common used genomic data files with a Python script or command line, interactively explore genomic data within Jupyter environment or web browser. Moreover, owing to the highly extensible API design, users can customize their own tracks without difficulty, which can greatly facilitate analytical, comparative genomic data visualization tasks.

## 1 Introduction

With the rapid development of next-generation sequencing technologies, more and more genomic assays have been developed for profiling the genome from various aspects, such as RNA expression[18], protein-DNA binding[20], chromatin accessibility[4] and 3D structure[16, 8]. By integrating data from these different kinds of assays or the so-called multiomics approach, biologists can comprehensively investigate genome dynamics during biological processes. This methodology has been successfully applied to many biological fields, such as neurological diseases[6], development of nervous system[22], virus infection[10, 5]. Data visualization, especially the genome track like plots, is crucial for exploring or demonstrating some local or global properties of the genome data.

Many visualization tools have been developed to meet these demands, and these tools can be classified into three categories: (1) Command-line plotting tool[17, 2], (2) Graphical User Interface(GUI) software[13], and (3) Web-based track browser[23, 14, 12]. Each of them has its own advantages and limitations for different situations; for example, command-line tools are convenient for bioinformaticians who prefer the command-line environment to quickly plot and check their results. GUI tools are friendly to people without programming skills. Web-based browsers enable users to share the visualization result with others over the internet. These kinds of tools work well for providing an overview of the input genomic data. However, during actual scientific research, we need more than just the basic view of the data. There are more needs for comparative and analytical data visualization; for example, to visualize the differential contact interaction(DCI) of two Hi-C contact matrices[3] or predicted chromatin loops on the matrix[21]. In most cases, bioinformaticians work in programmatic and interactive environments like RStudio, IPython console and Jupyter notebook to complete the data analysis, algorithm development and visualization tasks. However, there is a gap between the data analysis ecosystem and the existing genomic data visualization tools. Researchers spend on unnecessary stuff like file format conversion and environment switching. Therefore, a versatile tool that can fill this gap will significantly facilitate the genomics study.

To fill this gap, we developed CoolBox, a versatile toolkit for visual analysis of genomic data that combines advantages of existing tools, highly compatible with the Python scientific ecosystem, highly customizable, and easy to use with intuitive interface design and simple installation procedure. It can be used in different scenarios: (1) Python script or another python package for plotting and data fetching; (2) Shell as a command-line plotting tool; (3) Jupyter notebook environment for data fetching, plotting, and exploration; and (4) Web application for exploration and demonstration within a web browser.

## 2 Materials and methods

### 2.1 Requirement and installation

CoolBox is implemented with Python3; all dependencies can be installed and managed easily with conda(Anaconda package management tool) and the Bioconda channel[9]. CoolBox can be installed from the Bioconda channel using a single command line: “conda install -c bioconda coolbox”. Alternatively, users can utilize the latest features by installing from the source. CoolBox is developed and tested under Unix-based operating systems (Linux and macOS). Windows users can use it within Windows Subsystem for Linux(WSL) or Linux docker container.

### 2.2 Implementation

The plotting system of CoolBox is based on the matplotlib package. A part of the plotting code in the CoolBox is a fork from pyGenomeTracks[17] package. The data stored in bigWig, “.cool” and “.hic” file format are loaded through pybbi(https://github.com/nvictus/pybbi), cooler [1] and straw[7] package. Pairwise interaction data in BEDPE and Pairs format is indexed and randomly accessed using the pairix software (https://github.com/4dn-dcic/pairix). Other text-based genomic feature data format, like BED, GTF, and BedGraph is indexed and random accessed using the tabix[15] software. The widget panel in the GUI is implemented by using the ipywidgets package.

### 2.3 Availability

CoolBox is open-source under GPLv3 license at GitHub: https://github.com/GangCaoLab/CoolBox It can be downloaded from this site, or directly installed from the Bioconda channel. Detailed usage about API and CLI and various data visualization example is available in the online documentation: https://gangcaolab.github.io/CoolBox/index.html. An interactive online demonstration notebook about a small testing dataset is available on binder: https://mybinder.org/v2/gh/GangCaoLab/CoolBox/master?filepath=tests%2FTestRegion.ipynb.

## 3 Result and discussion

### 3.1 Flexible and user-friendly API and CLI for producing high-quality genome track plots

The interface of CoolBox includes an Application Programming Interface(API) for using it in Python script or Jupyter environment and a Command Line Interface(CLI) for using it in Shell. Its design is inspired by the popular R package ggplot2 [24]. It allows users to compose their figures with highly intuitive syntax. In CoolBox, users can use the “+” operator in Python or “add” command in Shell to compose low-level track elements to a higher-level figure. For example, they can compose track objects of various kinds of genomic data into a single frame and interactively review interested regions in genome browser with few lines of Python codes or Shell commands (Fig.1). Besides the 1-dimensional viewing mode supported by most other visualization tools, CoolBox supports a joint-view mode that enables users to visualize trans or cis-remote regions in a Hi-C contact matrix (Fig.2).

**Figure 1:**
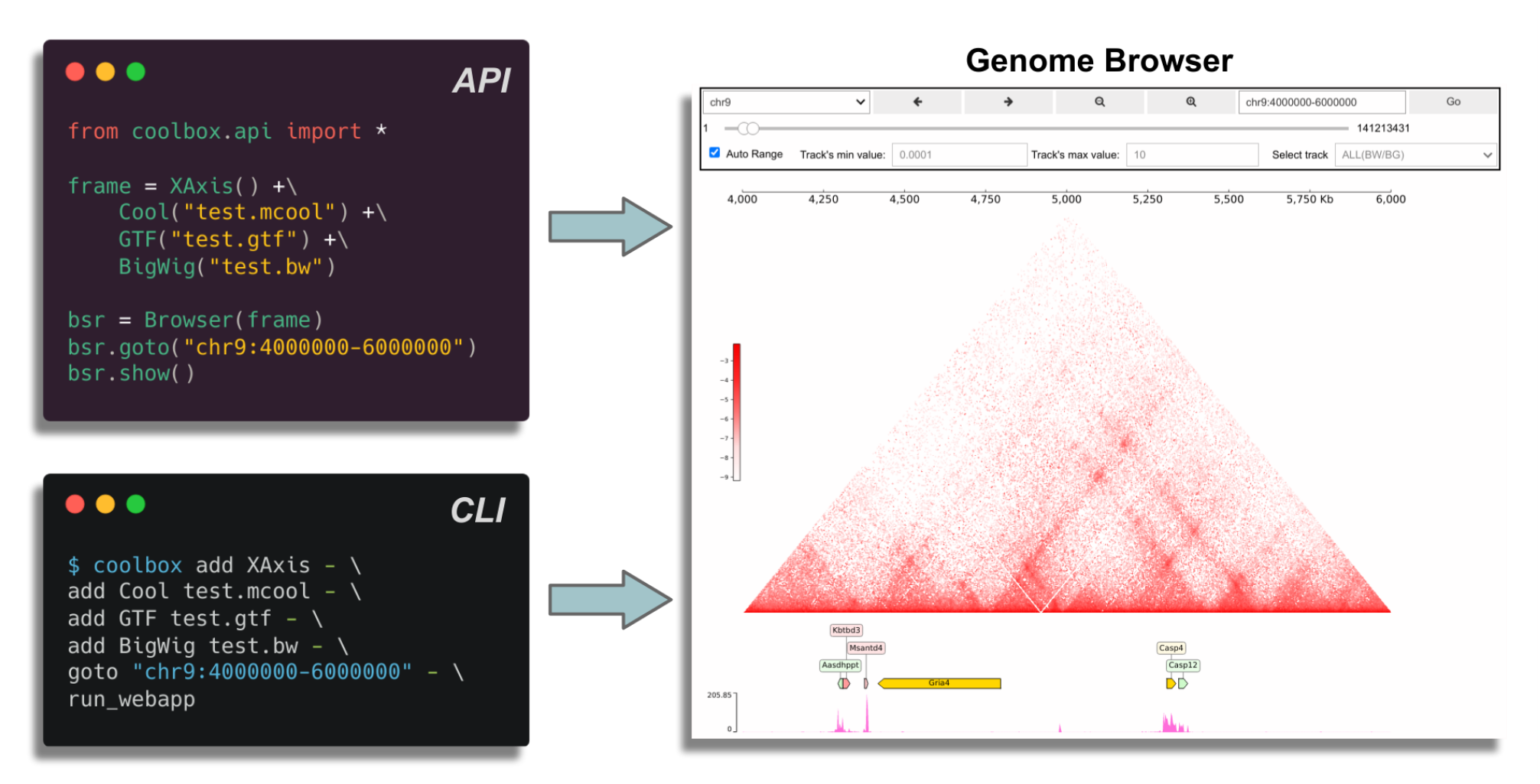
CoolBox has a clear and intuitive syntax to compose genome browser in both API and CLI mode. Inspired by the ggplot2 syntax, figures in CoolBox can be composed and adjusted(color, height, style etc.) from different tracks and features by using the ‘+’ operator in API or ‘add’ in CLI, almost every figure composed in the API mode has a paired CLI composing commands that produce identical figures. This design facilitates bioinformaticians that works usually in both environments.

**Figure 2:**
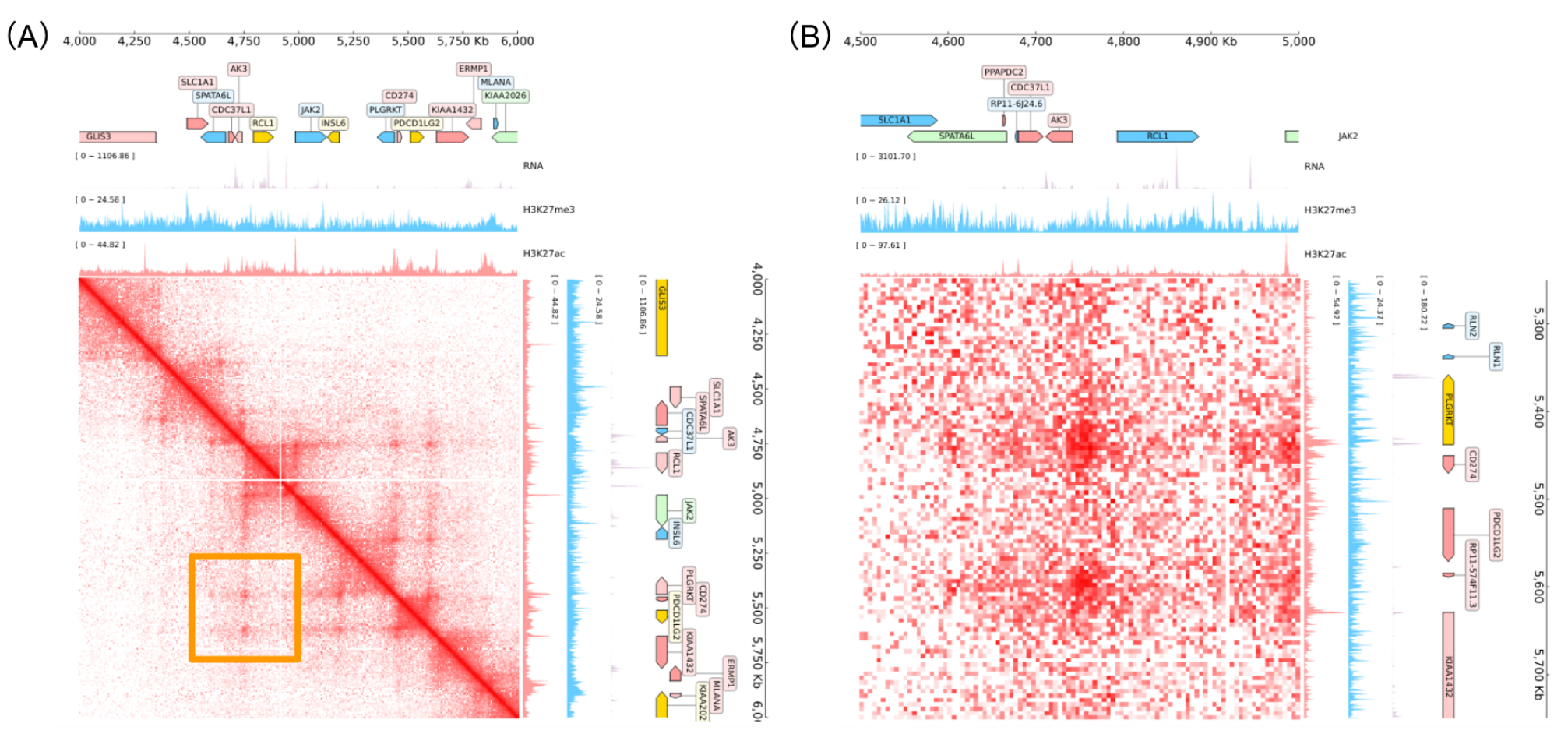
Joint(2d) view example, CoolBox can compose big figure which put frames around a center contact matrix. This allows to visualize the trans or cis remote(off-diagonal) contact matrix along with genome features. (A) Joint view on an on-diagonal region. (B) Joint view on an cis remote region, which shows the magnified detail of the orange box marked loop region that contains two chromatin loops in (A).

Most sets of commonly generated genomic assay data such as RNA-Seq, ChIP-Seq, ATAC-Seq, Hi-C, HiChIP[19] data which stored in bedgraph, bigwig[11], cool[1], .hic[7] file format (see Table.1) can be visualized in CoolBox by different kinds of tracks. Most tracks’ features (color, height, style, etc.) can also be configured the same way via the API or CLI. In the CoolBox plotting system, the plot contains not only a single layer. Users can put another layer (Coverage) upon the original plot to produce more comprehensive and high-quality figures. Moreover, the output figures can be conveniently saved in different kinds of image formats such as PNG, JPEG, PDF, and SVG.

**Table 1:** A part of CoolBox builtin tracks for visualizing different kinds of genomics data formats.

| Track type | File format | Description |
| --- | --- | --- |
| XAxis | None | Coordinate of the reference genome. |
| Spacer | None | For add vertical space between two tracks. |
| BigWig | .bigwig | Track for bigWig file, draw the histogram. |
| BedGraph | .bedgraph | Track for BedGraph file, draw the histogram. |
| BAM | .bam | BAM track for visualize the coverage or alignment. |
| BED | .bed | For visualization genome annotation, like refSeq genes and chromatin states. |
| GTF | .gtf | Track of GTF file, for visualize gene annotation. |
| Arcs | .pairs, .bedpe | Show the chromosome interactions get from ChIA-PET, HiChIP or Hi-C loop data. |
| HiCMat | .cool, .mcool, .hic | Show the chromosome contact matrix from Hi-C data. |
| Virtual4C | .cool, .mcool, .hic | Virtual 4C track, using Hi-C data to mimic 4C. |
| DiScore | .cool, .mcool, .hic | Directional index of Hi-C matrix for detecting TAD. |
| InsuScore | .cool, .mcool, .hic | Insulation score of Hi-C matrix for inferring TAD borders. |
| HiCDiff | .cool, .mcool, .hic | Show the difference between two contact matrix. |
| Selfish | .cool, .mcool, .hic | Apply the selfish algorithm[3] on two contact matrices to detect differential contact interactions. |
| SNP | .tsv | Track for show SNPs Manhattan plot. |

More details about how to use the API and CLI are available in the online documents.

### 3.2 Interactive exploration and reproducible analysis on genomic data

As shown in Fig.3, CoolBox provides a GUI for interactive data visualization, by which users can explore different genome regions by operating a simple widget panel and visualize the data within this region.

**Figure 3:**
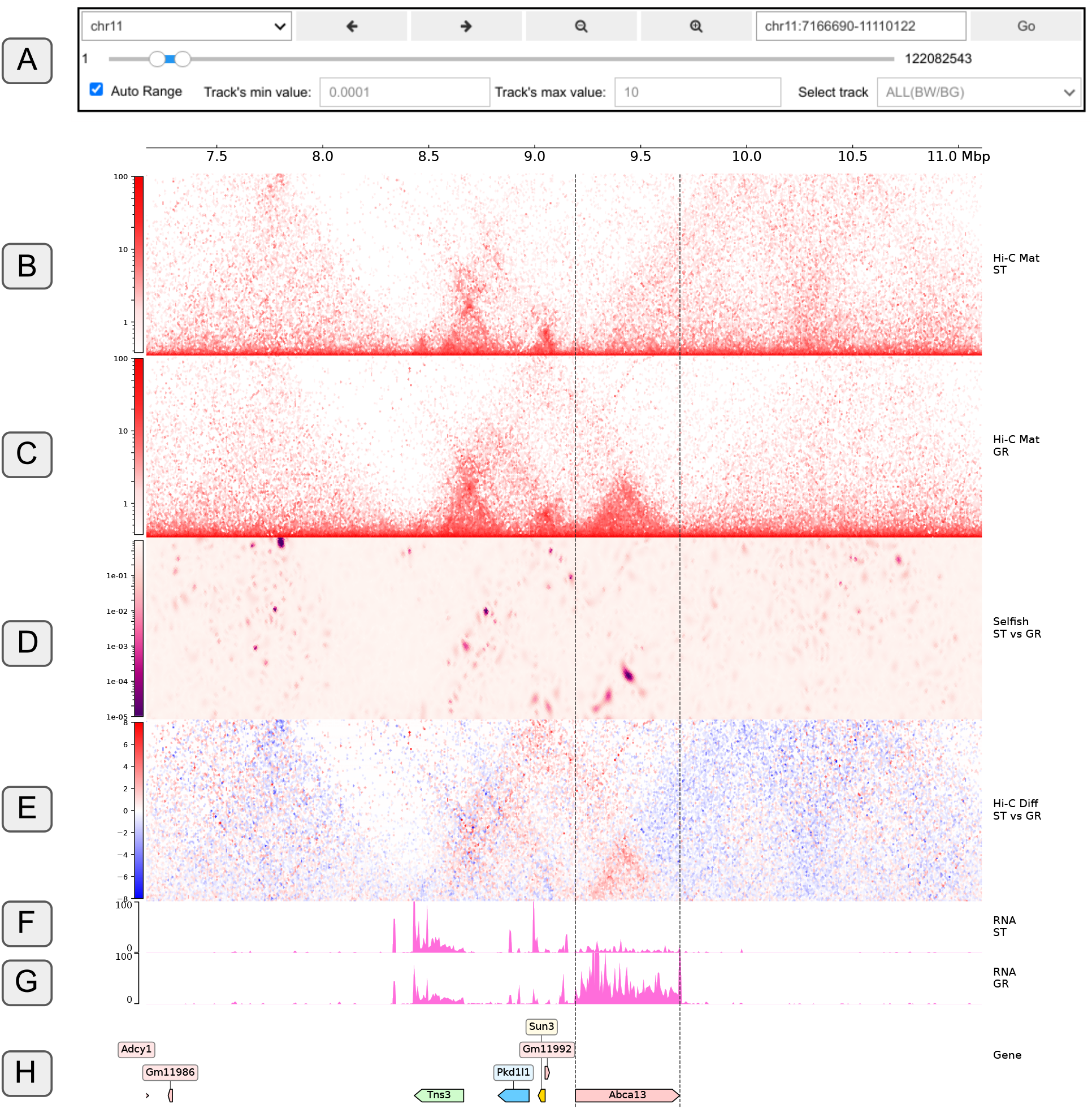
A CoolBox figure representing differential interactions of Hi-C contact matrices. Shown Hi-C and RNA-Seq data are produced from the process of hematopoietic differentiation[25]. Its clearly that there is a topological association domain(TAD) formation at the Abca13 gene region and its RNA expression is up-regulated at the same time after the differentiation. (A) The widgets panel of CoolBox browser, used for zooming, sliding, and locating the genome region. When moving to a new region, the figure draw bellow will be updated automatically. (B) Hi-C track of short-term hematopoietic stem cell(ST) shows the contact map of ST sample. The color bar indicates the normalized value of the contact map. (C) Hi-C track of granulocyte(GR). (D) Differential contact interaction(DCI) result of the Selfish algorithm[3] on ST and GR Hi-C contact map. The color bar indicates the q-value(BH adjusted p-value) produced from the DCI analysis. Darker color means this interaction has a lower q-value; that is to say, two contact maps are more diverse at this location. (E) Hi-C Diff track. It shows the difference between GR and ST’s z-score normalized contact matrices. The red region of the matrix indicates where GR has a more significant contact frequency compared to ST, and the opposite for blue areas. (F) BigWig track of ST RNA-Seq data, showing the RNA expression level of ST in this region. (G) BigWig track of GR RNA-Seq data. (H) A gene annotation track shows the corresponding genes within this genomic region.

Besides, the data and the figures are bound together by Python objects. In this way, users can get the precise data of each track within the current view of the genome region through the API. Such design facilitates comparative visualization and statistical analysis. CoolBox can also be used as a general genomic-file reading package. Data within a particular genome region can be retrieved in a short time, as almost all supported file formats can be indexed and randomly accessed.

Moreover, by leveraging the power of the Jupyter notebook, the visualization result and the entire process can be recorded in the notebook. It is convenient for sharing the visualization result and reproducing the whole analysis by other researchers.

### 3.3 A testing and visualizing framework for new algorithm development

Owing to the user-friendly and highly extensible API design, users can implement their custom tracks with no difficulty, thus enabling seamless cooperation in Python-based algorithm development and scientific research. The algorithm developer can check and visualize the intermediate result produced by their algorithm and adjust parameters simultaneously. Besides, because CoolBox uses an object-oriented programming paradigm in its design, users can reuse each track’s codes by inheritance, including data extraction and drawing-related functions. In most cases, users only need to write algorithm-related core parts, and the most tedious part including raw-data reading, preprocessing, and figure drawing are handed over to CoolBox through inheritance(implementation see method section). In this way, bioinformaticians can free themselves from those repetitive procedures and only focuses on the data post-processing.

We demonstrate these advantages by implementing a track that visualizes the outputs of the Peakachu algorithm[21], which is a RandomForest based method for detecting loops in the Hi-C contact matrix. As depicted in Fig.4, the main part of the whole track contains merely 20 lines of Python code. The data fetching and plotting functionality are fully reused by inheriting Cool/ArcsBase Track base class. Furthermore, the custom-defined track is empowered to be used in CLI, API, and browser mode in couple with other built-in tracks. More details of this example includes a reproducible code block and can be found in the online documentation.

**Figure 4:**
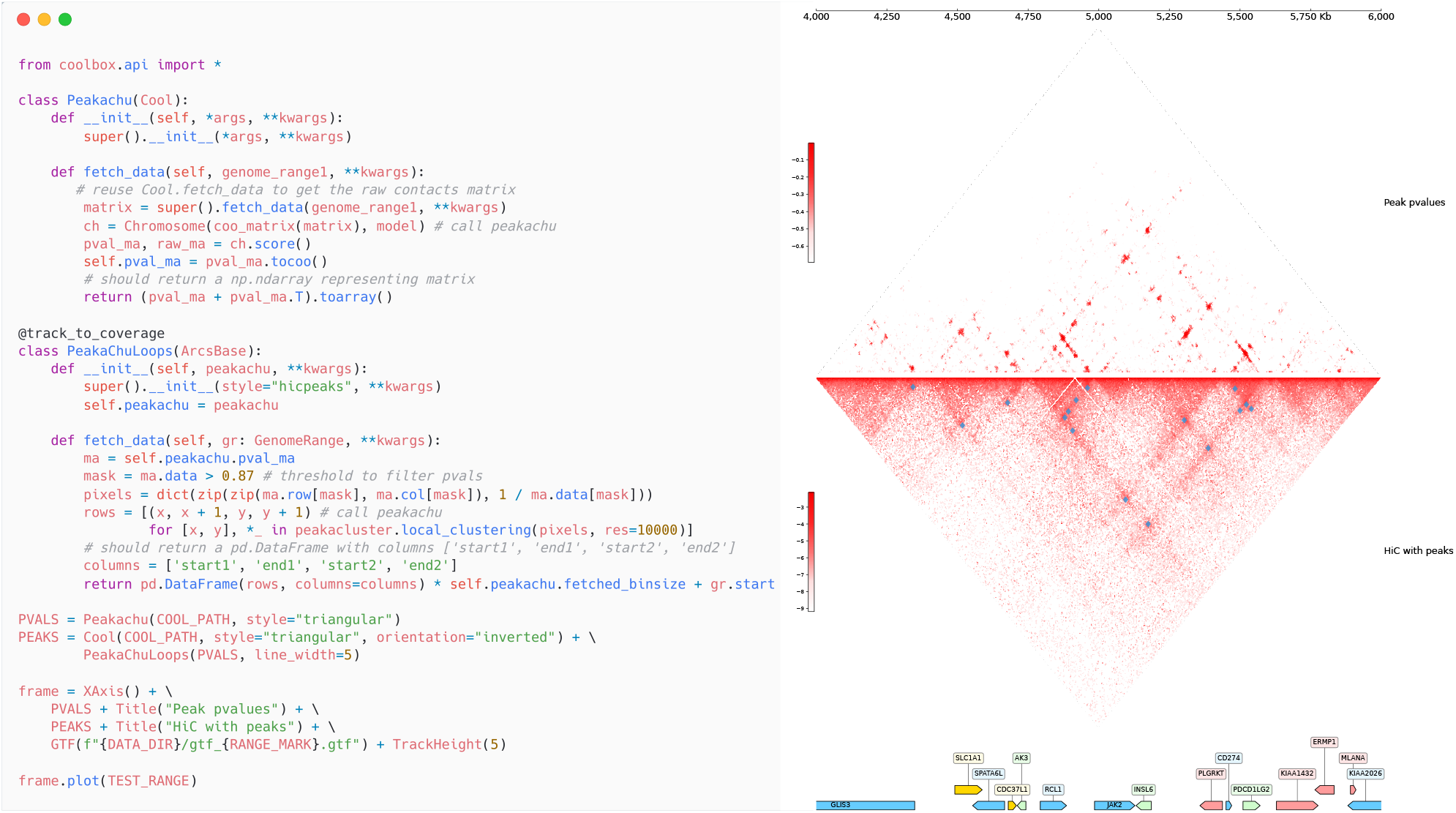
An example to define custom tracks that display Hi-C contact matrix along with peaks detected by Peakachu algorithm[21]. An example of peak prediction result is demonstrated in the right panel. The upper triangular matrix shows the peak p-value output by Peakachu algorithm. The predicted peaks drawn as blue squares upon the original matrix is shown in the lower triangular matrix. The Hi-C matrices and the peaks upon them will be automatically recomputed and updated after the change of genome region. The left panel is the full Runnable python codes used for generating the right panel. The custom track is implemented by following the same intuitive and clear design pattern as other built-in tracks: i.e., reusing the data fetching and plotting functionality as much as possible. For this figure, the functionality of fetching and plotting contact matrix with peaks are totally reused by inheriting Cool/ArcsBase track base class, and the rest of the codes merely calls the computing function of the peachachu package. After the track definition, we can see that the custom track is born to support being used in a ggplot2-like syntax with other tracks, and this capability is also valid in CLI and GUI mode.

### 3.4 Comparison with other existing visualization tools

As stated before, there is an urgent need for better visualization tools to accelerate the integration and mining of biological data. Therefore, more and more visualization tools have been developed in recent years. A comparison of features between CoolBox and these tools is listed in Table.2. Most of the visualization tools require a tedious installation process and are operated through the command line. Before visualization, the data needs to be preprocessed through specific steps, and then a static or interactive web interface is generated.

**Table 2:**
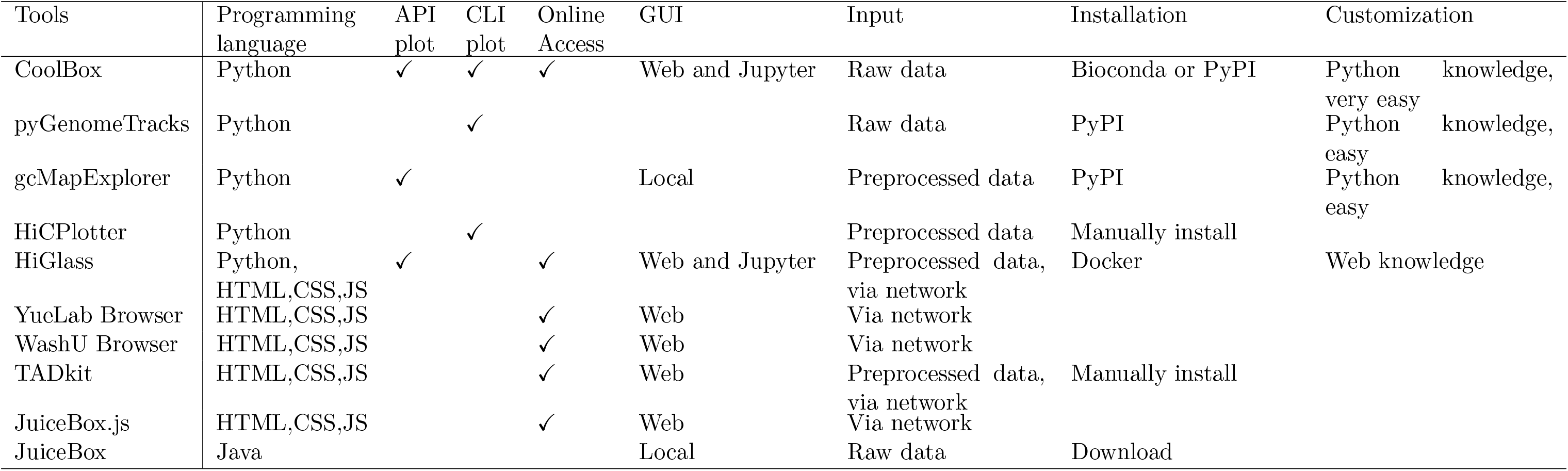
Summary of genomic visualization tools. The visualization and data processing of most visualization tools are dissociated, which is not convenient for bioinformaticians whose routine works rely on Python-based scientific computation ecosystem. Except for the CLI mode supported by most visualization tools, the API that the CoolBox has been used internally and exposed follows the same design idea as the CLI, making switching between these two modes with no pain. More importantly, the API in CoolBox combines computation and visualization, users can dynamically add different tracks or even custom tracks in the python notebook while processing raw data or developing new methods.

## 4 Conclusion

CoolBox is a versatile toolkit for the visualization and exploration of multiomics data in the Python ecosystem. It provides a user-friendly ggplot2-like syntax for composing various kinds of tracks in CLI, API, GUI and web browser mode. More importantly, it’s built on a highly extensible plotting system that allows users to implement their custom tracks without wasting time on data fetching and figure plotting procedures. Through the power of Jupyter notebook, it provides a convenient way for bioinformaticians to exploit this tool’s versatility for better personalized data manipulation and demonstration. It could also increase the reproducibility of genomic data visualization tasks as codes and figures are all organized into the same page.

